# NucleoNet and DropNet: Generalist deep learning models for instance segmentation of nuclei and lipid droplets from electron microscopy images

**DOI:** 10.64898/2026.04.02.713930

**Authors:** Abhishek Bhardwaj, Chris W. Dell, Melissa R. Mikolaj, Helen Spiers, Adam Harned, Balamurugan Kuppusamy, Peng Liu, Donglai Wei, Esta Sterneck, Kedar Narayan

**Author notes:** Corresponding Author (K.N.).

## Abstract

Automating cellular organelle segmentation is key to increasing the throughput in electron microscopy (EM) and volume EM (vEM) workflows. Deep learning (DL) has accelerated this process, but model development has predominately centered on mitochondria, partly because of a scarcity of suitable training datasets for other features. Here, we crowdsourced the manual step of labeling nuclei and lipid droplets (LDs) from complex cellular EM images and trained Panoptic DeepLab (PDL) models on these large, heterogenous annotated datasets as well as on publicly available vEM datasets. *NucleoNet* and *DropNet*, the resulting instance segmentation models for nuclei and LDs, respectively, yield high-quality results on varied benchmarks. We applied these models to quantify differences between 2D and 3D *in vitro* cancer models versus *in vivo* tumors, highlighting a path toward robust quantitation in EM. *NucleoNet* and *DropNet* are publicly available on our napari plugin, *empanada v1.2*, for easy point-and-click segmentation of 2D and 3D cellular EM images.

## Introduction

Electron microscopy (EM), and more recently, volume electron microscopy (vEM), have become indispensable tools for visualizing cellular ultrastructure at nanometer resolution in 2D and 3D, respectively (Collinson et al., 2023; Peddie et al., 2022; Varsano and Wolf, 2022). These approaches have been widely applied to study the spatial organization and structural properties of organelles such as mitochondria, nuclei, endoplasmic reticulum, and LDs (Briggman and Bock, 2012; Peddie et al., 2022; Titze and Genoud, 2016; Xu et al., 2017), often requiring lengthy and often complicated experimental procedures (Genoud et al., 2018; Hua et al., 2015). Moreover, the resulting image datasets are vast, in both volume and complexity; these attributes present a major challenge for downstream data analysis. This is particularly acute for image segmentation and quantitative assessments of organelle morphology, where few, if any, automated solutions are available (Kubota et al., 2018; Shapson-Coe et al., 2024). Indeed, manual segmentation of organelles from EM images remains the gold standard for high accuracy annotations even though it is notoriously time-consuming and labor-intensive.

This challenge has driven the development of computational approaches designed to standardize, accelerate, and automate segmentation; these efforts have achieved some success and notable recent advances (Archit et al., 2025; Belevich and Jokitalo, 2021; Chen et al., 2017; Falk et al., 2018; Haberl et al., 2018; Jiang et al., 2025). However, most organelle segmentation tools have focused on mitochondria (Berning et al., 2015; Conrad and Narayan, 2023; Jiang et al., 2025; Wei et al., 2020), possibly because of their biological significance, relative abundance, and ease of visual detection, while other crucial structures like nuclei and LDs remain comparatively underexplored. This is in contrast with fluorescence microscopy, where nuclear segmentation is facilitated by easily resolved instances imaged at high signal-to-noise ratios, resulting in high-quality models (Achard et al., 2025; Riendeau et al., 2025; Sankaranarayanan et al., 2025), whereas smaller and closely packed organelles are more challenging to correctly resolve and segment. To our knowledge, no ready-to-use tools currently exist for automated instance segmentation of nuclei and LDs in 2D and 3D EM images. A commendable effort by Jones and colleagues resulted in a model for semantic segmentation of the nuclear envelope (Spiers et al., 2021a) https://github.com/FrancisCrickInstitute/Etch-a-Cell-Nuclear-Envelope. The model is based on a U-net architecture trained on images derived from a single, massive serial block face SEM (SBF-SEM) volume comprising several HeLa cells. In our hands the model required significant computational expertise to set up and we were unable to run inference on 2D images, underscoring to us the importance of usability of software tools. In this paper: 1. we train and test/evaluate models for segmentation of nuclei and LDs specifically to be generalist, ensuring broad applicability across EM datasets, and 2. we incorporate these models into an easy-to-use GUI, enabling accessible, point-and-click utility for the wider community.

The successful application of deep learning (DL) models depends on the availability of widely annotated, high-quality datasets that are relevant, information-rich, non-redundant, and heterogenous (Caicedo et al., 2019; Meijering, 2020; Moen et al., 2019). For mitochondria, extensive unlabeled and labeled datasets such as MitoEM, CEM500K and 1.5M, and CEM-MitoLab (Conrad and Narayan, 2021; Wei et al., 2020) enable the development of powerful and generalist models. In contrast, datasets with annotations for nuclei and LDs remain scarce. Two notable exceptions are: 1. The MICrONS dataset (Zhang et al., 2025), a petabyte-scale serial section TEM image volume of the mouse visual cortex with a sub-volume of semantically segmented nuclei, and 2. A terabyte-scale SBF-SEM of whole-body volume of the nereid *Platynereis dumerilii* with good but not perfect instance segmentation of nuclei (Vergara et al., 2021a); we leverage both in our model training. Models trained solely on a single homogenous 3D dataset (Spiers et al., 2021a; Zhang et al., 2025) likely generalize poorly because they lack diverse cell types, tissues, and are limited to a single sample preparation and imaging protocol (Conrad and Narayan, 2021). One promising strategy to overcome this annotation homogeneity is crowdsourcing of diverse datasets. In the biomedical imaging community, crowdsourcing has been successfully applied to annotate light microscopy images, histopathology slides, and even EM images (Bafti et al., 2021; Carreras et al., 2015; Spiers et al., 2021). Platforms like Zooniverse allow scientists and citizens to contribute to large annotation efforts by distributing the annotation workload, thus helping generate large training datasets essential for deep learning(Simpson et al., 2014; Smith et al., 2023). But claims of superior model performance must be supported by objective metrics. A gap between published results and post-publication model performance points to a lack of accessible and challenging benchmark datasets in the EM field; this contrasts with other computer vision areas, where such benchmark resources are widely adopted and highly valued (Guo et al., 2014; Salari et al., 2022). Finally, models must be usable by non-experts. Regardless how sophisticated and powerful a model is, it will be underutilized unless it is compatible with various computational environments and is straightforward to set up and run, especially for researchers in resource- and expertise-limited environments, seeking an easy-to-use solution for their segmentation task (Kniesel et al., 2025; Meyer et al., 2025).

To address both the annotation bottleneck and the need for usable, generalist models, we developed crowdsourced annotation pipelines tailored for complex EM datasets and used these curated annotations to train segmentation models for nuclei and LDs. Employing the Panoptic DeepLab (PDL) architecture, which enables efficient and accurate segmentation in both semantic and instance domains, we created two models: *NucleoNet* for nuclear segmentation and *DropNet* for lipid droplet segmentation. These models were trained on diverse EM datasets and validated on unseen benchmarks, spanning cultured cells, tumor samples, and tissues, to demonstrate their generalizability and robustness. Both models operate in two dimensions, enabling relatively rapid and resource-efficient segmentation of any 2D EM image, and they can also be used to segment 3D features by running inference in the imaging and orthogonal planes within image volumes, as previously described (Conrad et al., 2020). We were able to compare *NucleoNet* with two recently released models, CellSAM (Marks et al., 2025)and nnU-net (Isensee et al., 2020a), and we show that it far outperforms CellSAM and is comparable to nnU-net, while having the advantage of being easy to use. We use *NucleoNet* and *DropNet* to generate large numbers of high-quality segmentation label maps in various *in vitro* cancer models, enabling statistically significant comparisons of various morphological metrics against an actual tumor sample; more biological details are included in a recently published study (Balamurugan et al., 2026). Additionally, we also release a new version of our napari plugin, *empanada*, which now comes pre-packaged with *NucleoNet* and *DropNet* (as well as our original models *MitoNet* and *MitoNet_mini).* The plugin features versatile modules for proofreading, model fine-tuning, etc, along with thorough documentation (https://empanada.readthedocs.io/en/latest/index.html); with >110,000 downloads, it is popular within the vEM community.

## Results

Training datasets are critical to model performance (Conrad and Narayan, 2021; Lu et al., 2024). In previous work, we used crowdsourced annotation strategies to create massive datasets of instance-labeled mitochondria to create *MitoNet* (Conrad and Narayan, 2023). Here, different constraints demanded two distinct strategies to create DL segmentation models for nuclei and LDs.

### NucleoNet

We collated EM images from two sources: an internal corpus comprising a combination of mostly unpublished EM images from focused ion beam scanning electron microscopy (FIB-SEM), SEM array tomography (AT) and conventional transmission EM (TEM); and an external corpus derived from publicly available repositories such as EMPIAR (Iudin et al., 2023), OpenOrganelle (Heinrich et al., 2021) and nanotomy (de Boer et al., 2020) which also included SBF-SEM images. In total, we gathered approximately 4.8 x 10^6^ 2D images from external sources and 2.5 x 10^6^ 2D images from internal sources, including images sampled from orthogonal views in 3D datasets. Near-identical and non-informative images were removed to create a corpus of approximately 450,000 patches (1024 x 1024 pixels; see Materials and Methods), with an internal-to-external source ratio of roughly 2:1. These patches covered a variety of organisms, organs, and experimental systems, capturing a broad range of nuclear shapes and staining patterns **(Supp Fig S1)**. Based on biological and image metadata, these patches were enriched for presence of nuclei and further screened manually by experts for human annotation suitability based on uniqueness and difficulty – patches deemed to be too ambiguous or too similar were discarded. From this set of ∼70,000 heterogenous, information-rich and challenging images, random image subsets (∼500 image patches per batch) were uploaded to the crowdsource annotation platform Zooniverse.

High-school students were trained over several remote sessions to create accurate label maps outlining nuclei in the uploaded images. Annotations were performed entirely remotely on the Zooniverse web browser GUI, and participants were asked to complete manual annotations of 500 images per week. The Zooniverse platform provided modules to share examples of good and poor annotations, tips for good practices, relevant biological and imaging background, and a chat channel. Each image was annotated by five different students to enhance segmentation quality, with pixel-level accuracy encouraged. In parallel, an expert annotated fifty images per batch to provide putative ground truth (GT), which was used to evaluate accuracy scores for all users. All student and expert annotations were pooled and aggregated into a single consensus label map annotation for each image, which was proofread by two experts at the end of every weekly session to create batches of GT training data **(Fig 1A)**. Typically, 20-25 students participated each week, and we observed a weak inverse correlation of segmentation efficiency and accuracy against difficulty and novelty of the image sets **(Fig 1B)**. In total, 32,745 crowdsourced annotations were generated and proof-read by experts, corresponding to 6,549 high-quality annotated image patches.

**Figure 1.**
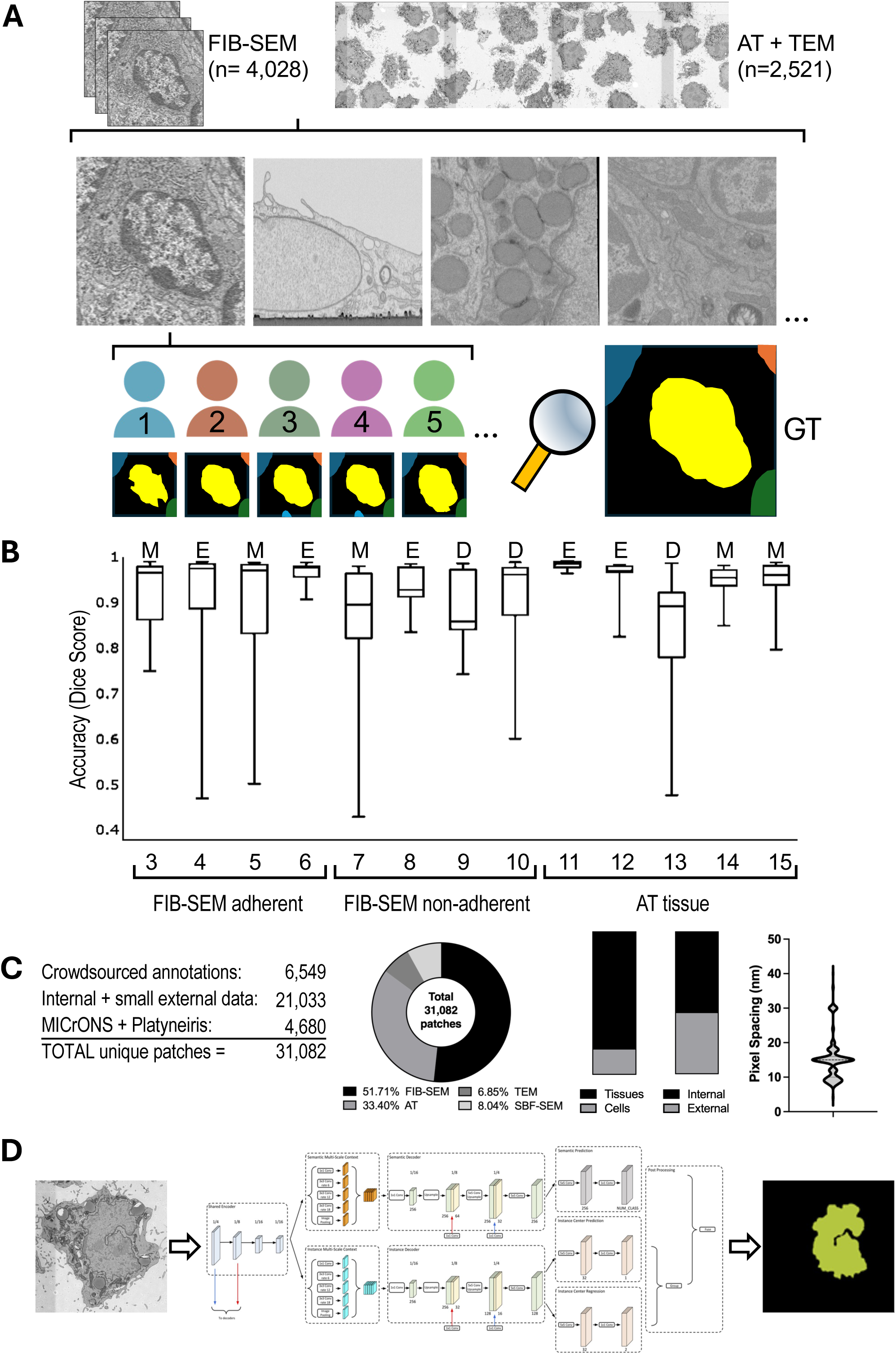
Training dataset generation and strategy for *NucleoNet*. (A) Image patches (512 x 512 pixels) from FIB-SEM, AT, and TEM datasets were distributed at random and each image annotated by five annotators; a final ground truth (GT) was generated by combining the annotations and expert proofreading. (B) Annotator segmentation accuracies for dataset batches, with image batches visually graded as E, easy; M, medium; D, difficult. Box = mean, std dev; whiskers = min, max. Datasets 1 and 2 were trials and discarded. (C) Breakdown of *NucleoNet* training dataset from left to right by annotation source, imaging approach, cell context, internal versus external, and image pixel size. (D) Schematic of Panotopic DeepLab (PDL) architecture used for model training.

In parallel, we also downloaded roughly 710,000 and 1.25 x 10^6^ patches and their paired nuclear segmentation from the MICrONS and platyneiris datasets (Vergara et al., 2021a; Zhang et al., 2025), respectively. We tuned the fractional representation of these two sources in the training dataset and found that a 15% total contribution optimized model performance. Thus, 2,130 serial section TEM (ssTEM) images (MICrONS) and 2,500 SBF-SEM annotated images were combined with 21,033 images containing pre-segmented nuclei (from internal projects and smaller publicly available datasets) and 6,549 crowdsourced annotations to yield 31,082 labeled images for training. Grouped by source or data provenance, these comprised 17,746 internal and 13,386 external images. Together, this was a heterogenous training dataset based on source, imaging approach, cell context (in vivo/tissue or in vitro/cells) and pixel sampling **(Fig 1C, top)**.

The *empanada* library, including the PDL model architecture (Cheng et al., 2019a) was used for model training, following a strategy like that previously used for *MitoNet* **(Fig 1C, bottom)**. Briefly, the model was pre-trained on CEM1.5M, a dataset previously shown to be beneficial for model generalization (Conrad and Narayan, 2023). The encoder was initialized with these pre-trained weights, and a progressive unfreezing strategy was used to fine-tune encoder layers. Final models were selected based on the highest average precision achieved on a held-out validation set, as detailed in Materials and Methods. The PDL architecture yields both semantic predictions and X/Y offsets, which can be combined to generate instance predictions. Both *NucleoNet* and *DropNet* are 2D models, allowing fast performance in resource-limited environments. For 3D data, segmentation is performed by running inference in orthogonal planes (xy, xz and yz), which is a module accessible in the plugin UI.

The base *NucleoNet* model was trained to produce reliable segmentation results for nuclei in a variety of cellular contexts. For example, in volume EM images of a tumor biopsy specimen, *NucleoNet* accurately segmented widely heterogenous nuclei, including those with deeply invaginated membranes **(Fig 2A, arrowhead)**. Nevertheless, split and merge errors occasionally occurred, with false instance splits dominating in some images **(Fig 2A, boxed)**; these errors can be easily corrected using the split and merge modules in the *empanada* plugin. *NucleoNet* also generated good segmentation results in 3D using the “orthoplane” module in the *empanada* napari plugin, with weaknesses in some planes often compensated by strong predictions in others. We observed some errors **(Fig 2B, boxed)** appearing as a cross-hatched pattern in small sub-volumes, likely because of borderline predictions crossing the threshold intermittently; these are also trivial to smooth out during proof-reading. As with all current models, false negatives (FN), false positives (FP) occur; occasionally patches near the centers of nuclei that do not contain nuclear membranes may be missed. Again, these errors can be mitigated by adjusting confidence and other inference parameters; in the *empanada* workflow downsampling by 2 or 4 to yield a resolution in the 15-30 nm (see Discussion) along with a center distance above 30 works well. In case of spurious false positives, a 0.6 inference confidence threshold gives better results, and conversely, a 0.3 inference confidence reduces large false negatives in our hands.

**Figure 2.**
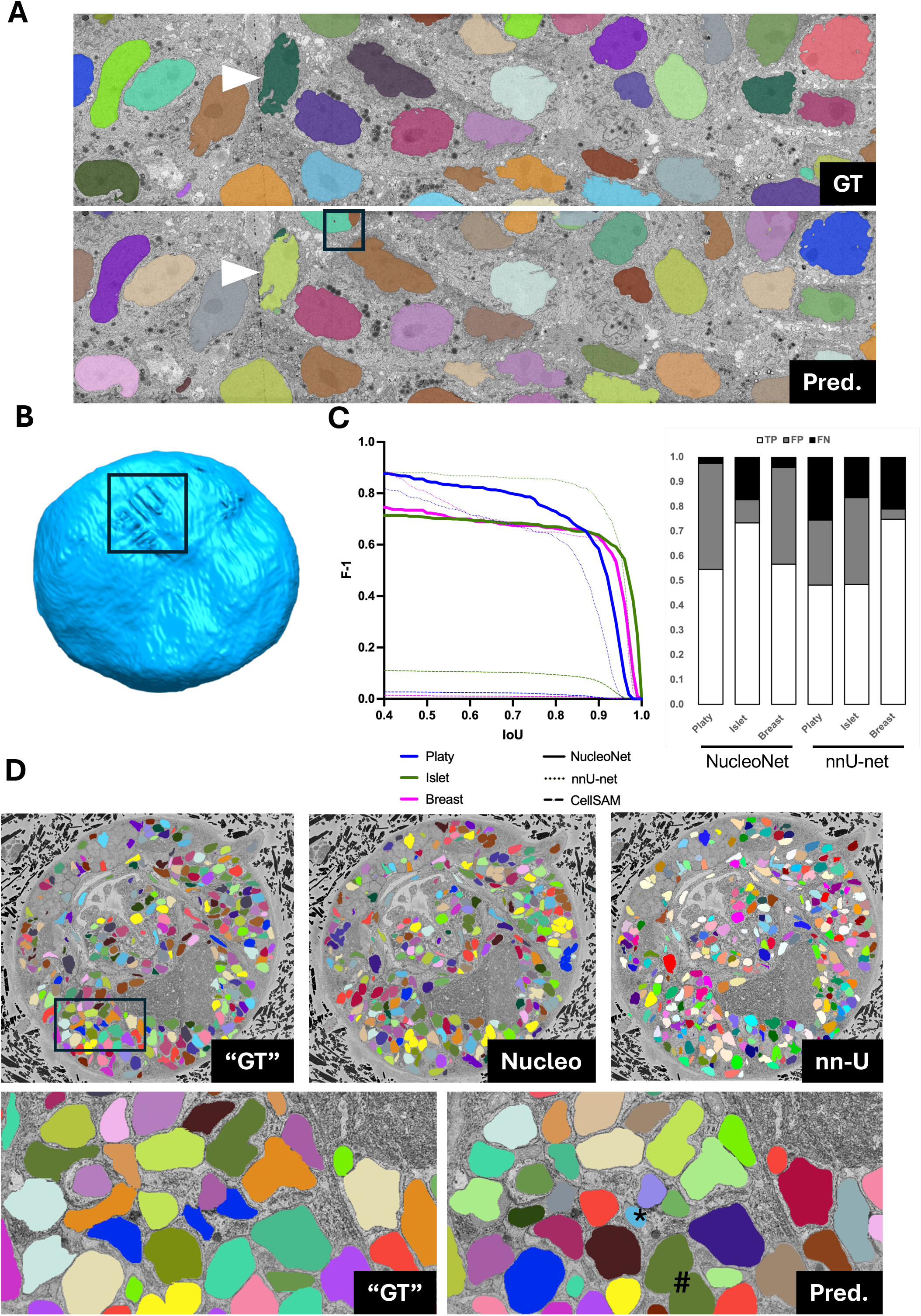
Evaluation of *NucleoNet* performance. (A) Representative 2D patch of a breast cancer-derived spheroid imaged by array tomography, showing variable and convoluted nuclei (arrowhead), overlaid with ground truth (GT) instance segmentations (top) and *NucleoNet* model prediction (bottom). Box highlight nuclei regions with split errors. The Horizonal Field Width for the snapshots are 225 μm. (B) Representative 3D volume segmentation produced by running *NucleoNet* across xy, xz, and yz planes. Box indicate segmentation errors arising from prediction mismatches. (C) Plot of F1 score versus intersection over union (IoU) threshold, showing segmentation accuracy over increasing overlap stringency between predicted and ground-truth nuclei (left). *NucleoNet* results are bolded, nnU-net dotted, and CellSAM dashed lines. Benchmarks tested are *Platynereis dumerilii* (Platy, blue), Rat islet of Langerhans (Islet, green) and berast cancer derived spheroid (Breast, pink). The corresponding fractional bar graph illustrates false negatives (FN, black), false positives (FP, grey), and true positives (TP, white) across the three benchmarks. Low scores from CellSAM was excluded for clarity. (D) Top: Large area of nereid *Platynereis dumerilii* putative ground truth segmentation (“GT, left) compared to *NucleoNet* model prediction before (Nucleo, middle) and nnU-net (nn-U). Bottom: Boxed sub-area of ground truth and corrected labels, showing occasional merge errors (#) and over-erosion (*). Horizonal Field Widths are 250 μm and 75μm for the overview and cropped images, respectively.

We tested the model against new and challenging benchmarks, and also against newer model architectures. Unfortunately, unlike benchmarks available for mitochondria (Guay et al., 2021; Lucchi et al., 2012, 10.6019/EMPIAR-10982) there are few benchmarks for nuclei, especially to test generality. Consequently, we selected vEM images from diverse biological samples, specifically choosing those with high variability, multiple nuclear shapes per image, varied sample preparation and different imaging approaches. Specifically, we chose the following: 1. A random section from the publicly available Serial Block Face Electron Microscopy Volume (https://www.ebi.ac.uk/empiar/EMPIAR-10365/) of a model organism, platynereis dumerilii (Vergara et al., 2021b), 2. A publicly available section of a rat islet of Langerhans imaged by TEM (https://nanotomy.org/) (Ravelli et al., 2013), and 3. A recently published Array Tomography dataset of a section through a human breast cancer derived spheroid(Balamurugan et al., 2026). *NucleoNet* performed reasonably well and roughly comparable F1 vs IoU curves against benchmarks **(Fig 2C)**, with F1@0.50 results ranging from 0.633 to 0.818 **(Table 1)**, which we interpret as a marker for model robustness to image variability. We acknowledge that images of Platy were present in the training dataset, but note that only ∼0.038% of this large dataset was included in training and that the accompanying nuclear segmentations were not pixel perfect; it is unlikely to have an undue influence on model performance, which was in line with the other benchmarks. *NucleoNet* predictions were high-quality **(Fig 2D, top middle)** with a mean pixel level IoU of 0.961 and mean instance IoU of 0.873 **(Table 2, left)**. However, limitations and compromises with the model resulted in two errors – a number of small false positives, and some expected split/merge errors with a bias toward false merges. This resulted in low precision and F-1 scores of 0.5759 and 0.6876, respectively. Importantly, when we executed the following steps: filter all instances below 500 pixels, erode by 40 pixels, connected components, dilate 40 pixels (altogether a few clicks in the napari GUI), precision and F-1 scores increased to 0.9367 and 0.8929, respectively. FPs fell by 90% as well **(Table 2, boxed)**, and the results were visually improved. Most tightly apposed nuclei were correctly separated **(Fig 2D, bottom)** but a few stayed merged (#) and some nuclei were too aggressively eroded (*), so final proofreading is required. These results are with the base model, so further improvement is predicted with fine-tuning.

**Table 1.** Evaluation of *NucleoNet* model performance against five varied benchmarks. TP, True Positive; FP, False Positive; FN, False Negative.

**Table 2.** Pixel-level and Instance-level metrics of NucleoNet predictions on a random slice of the Platyneiris dataset, either directly after running inference (left two columns), or after running the following steps available in the napari plugin: filter out all instances below 500 pixels, erode by 40 pixels, run connected components (to establish new instances), dilate by 40 pixes.

We also compared *NucleoNet*, which is a relatively older Panoptic DeepLab model (Cheng et al., 2019b), against two newer frameworks: 1. nnU-Net (Isensee et al., 2020b), a popular self-configuring architecture with excellent results on biomedical images, which was trained on a high-quality and heteroengous volume EM dataset (see details in Methods), and 2. CellSAM (Marks et al., 2025), a recently released foundational model tuned to cell segmentation across a variety of cell types and imaging approaches. In line with our expectations, nnU-Net results, being a larger model trained on EM images, gave good nuclear predictions; after running a watershed algorithm to convert semantic to instance segmentations and running a size filter, the outputs were comparable to zero-shot *NucleoNet*, with surprisingly good performance on the Islet benchmark **(Fig 2C, dotted line)**. The breakdown of FP and FN at IoU 0.75 showed that *NucleoNet* tended to yield fewer FNs, but visually there was little to choose between the models. On the other hand, CellSAM gave very poor results both statistically **(Fig 2C, dashed line)** and visually (data not shown). While current models are far from perfect, several aspects of *NucleoNet* render it favorable: it is tuned to EM images (unlike CellSAM), and it is available as an option in *empanada* for easy use by non-experts (unlike nnU-Net). Further, we see large improvements in segmentation with simple post-processing steps, and of course *NucleoNet* can also be fine-tuned to specific cellular EM data. Combined with cleanup tools also available in *empanada,* this is an immediately useful deep learning model for instance segmentation of nuclei in varied EM and vEM image data. Future models will perform undoubtedly perform better with these and newer benchmarks and with challenging biological samples.

### DropNet

LDs are less prevalent in available cellular EM datasets and tend to be small, spherical, and exhibit characteristic staining patterns with conventional EM staining protocols **(Supp Fig S2)**. To address this variability, we categorized LDs into four visual classes: 1. droplets of varying contrast with a dark outline, 2. droplets of varying contrast without a dark outline, 3. droplets with an unstained center, and 4. droplets displaying artifacts such as dots, cracks, aspherical shapes, or unusual staining patterns **(Fig 3A)**. Importantly, osmophilic granules that closely resemble certain LDs in terms of size, shape, and staining, such as those in acinar cells of salivary and pancreatic tissues, were classified as true positives (TP) **(Fig 3B, top)**. On the other hand, glucagon, insulin, and somatostatin containing granules from alpha, beta, and delta cells of the islet of Langerhans, respectively, were classified as true negatives (TN) **(Fig 3B, bottom)**. This distinction is important for downstream use, as the base model will segment secretory acinar granules and ignore the other pancreatic granules. Therefore, further fine-tuning of *DropNet* to exclude acinar granule segmentation or conversely, to include insulin granule segmentation, is not advisable.

**Figure 3.**
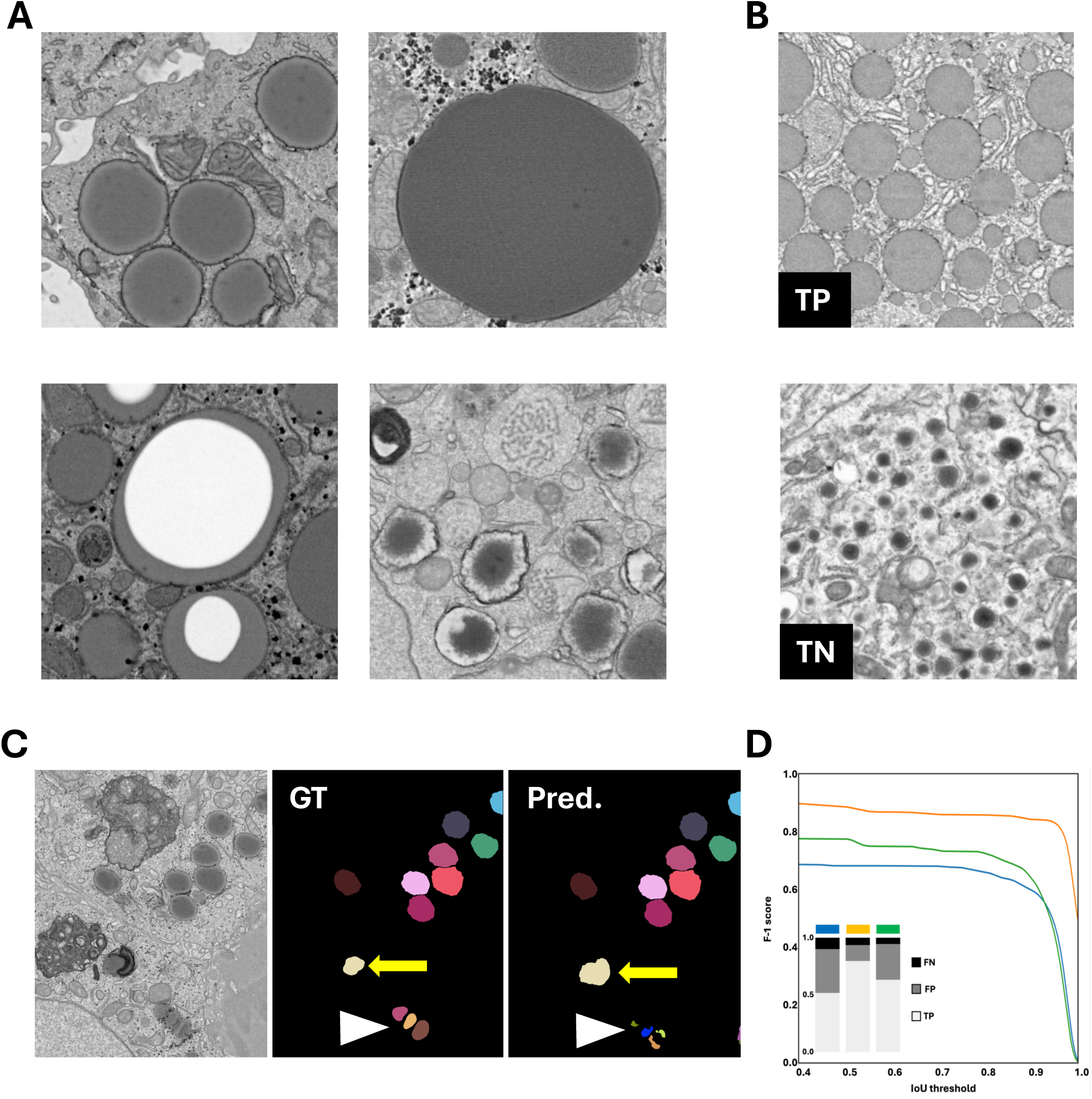
Lipid droplet classes and *DropNet* performance. (A) Examples of lipid droplet classes: variable staining with distinct dark outline (top, left), lack of outline (top, right), a central “hole” caused by sample preparation artifacts (bottom, left), and irregularly shaped or unevenly stained (bottom, right). (B) Round, uniformly stained salivary or acinar granules (top) were considered true positives (TP), whereas distinct, characteristically shaped insulin granules (bottom) were classified as true negatives (TN). Horizonal Field Widths for snapshots in A and B are 4 μm. (C) Representative ground truth segmentation (middle) and the corresponding *DropNet* model predictions (right), with false negatives (white arrowhead) and a false positive (yellow arrow). The original EM data is shown in the left panel. (D) Plot of F1 score versus IoU threshold with accompanying fractional bar graph showing the proportions of false negatives (FN, black), false positives (FP, dark grey), and true positives (TP, light grey) across three distinct benchmarks: blue – fatty liver; yellow – pancreas; green – breast cancer.

Overall, we were able to collate 1,792 TP image patches of 512 x 512 pixels across the identified LD classes. These patches were manually picked from FIB-SEM, AT, and TEM datasets, with each patch containing between 1 and 50 LDs, as these structures tend to cluster in the cytoplasm. Identifying lipid-rich patches in publicly available datasets proved challenging, apart from nanotomy.org; thus, most patches were sourced from internal datasets, primarily from liver, kidney, breast, and pancreatic tissues, as well as tumor cell lines and model organisms. LDs were manually segmented using thresholding and cleanups in 3DSlicer, and 250 TN images containing targeted non-LD granule structures were added to the TP image set. Given that the number of training patches was limited, we opted to fine-tune an existing internal variant of the *MitoNet* model with these patches, with all layers fine-tuned to the new data (see Materials and Methods). As a result, *DropNet* is less robust than *NucleoNet*, constrained by the size and heterogeneity of the training dataset. The model was tested against three previously unseen benchmarks (indeed holding out these datasets further shrank the training corpus) and performed well. Most instances of LDs were correctly identified and segmented, but unusual droplets with staining artifacts or close proximity to other features resulted in FP and FN errors **(Fig 3C)**. Even when the droplets were tightly clustered, the model correctly separated individual instances, rarely requiring split/merge corrections. F1@0.75 scores on the three benchmarks were 0.770, 0.840 and 0.911, with errors predominantly from FPs **(Fig 3D**, **Table 3)**. Importantly, islet cell granules from internal and external datasets were completely ignored by the model, while acinar granules were correctly segmented (data not shown).

**Table 3.** Evaluation of *DropNet* model performance against three varied benchmarks. TP, True Positive; FP, False Positive; FN, False Negative.

### Application to tumor models

*NucleoNet* and *DropNet* were used to test the relevance of various established and novel *in vitro* tumor models in comparison to an excised tumor specimen of the same cell type, based on ultrastructural characteristics of nuclei and LDs. Detailed biological findings are not presented here as a separate manuscript covers this topic (Balamurugan et al., 2026). Instead, we highlight the qualitative and quantitative results enabled by these models for accurate and precise segmentation of these organelles in disparate model systems. A xenograft breast cancer tumor derived from SUM149 cells was designated as the “tumor” control, while four *in vitro* models derived from the same cell line: “Adherent” (cells were plated in culture, representing a 2D model), “Suspension” (cells were grown in a non-adherent manner and pelleted before fixation), “Spheroid” (cells were allowed to aggregate into a sphere-like clump of cells) and “Emboli” (aggregation was accompanied by increased media viscosity and mechanical rocking to recapitulate blood flow) were also analyzed **(Fig S3A)**. Cells from each condition were fixed and processed for electron microscopy; AT data is presented here to showcase the power of these models to generate many segmentations at a single click. We note that identical models were used for all samples without further fine-tuning or manual adjustments. The only interventions were corrections of occasional split/merge errors after running inference; these could be executed with a few clicks in the appropriate module in *empanada*. For accurate quantitation, instances touching image borders were excluded from analysis.

Representative segmentation outputs are shown in **Fig 4**; note all nucleus and LD instances cut off at image edges were segmented but deleted for downstream analysis and therefore are not visible. Adherent cells exhibited the most complex nuclei **(Fig 4A, top)**, yet *NucleoNet* accurately segmented all instances, including nuclei with deep invaginations **(Fig 4A, top and middle boxed; 4B, left)**. Results were optimal when the images, which were collected at 10 nm pixel size, were downsampled by 4. These cells contained variable numbers of LDs, but the staining was relatively straightforward, allowing *DropNet* to generate correct segmentations with minimal or no human intervention, even when LDs were tightly clustered **(Fig 4A top and bottom boxed; 4B, right)**. For this manuscript, equally sized image strips of cells from each growth condition, as well as the tumor, were cropped to obtain comparable numbers of cells per group, with the more sparsely distributed adherent cell group at the lower bound. Nuclei instances (excluding edge cases) from the groups were as follows: adherent, n = 18; emboli, n = 32; spheroid, n = 25; suspension, n = 43; tumor, n = 32. The LDs from the groups were: adherent, n = 73; emboli, n = 319; spheroid, n = 187; suspension, n = 258; tumor, n = 91. From these instance segmentations, we generated basic morphometric measurements using the RegionProps plugin in napari. We plotted area, solidity and eccentricity of the nuclei **(Fig 4C, top)** and LDs **(Fig 4C, bottom)** and performed parametric unpaired t-tests with Welch’s correction, comparing each *in vitro* models against the tumor group. These analyses revealed that the different *in vitro* models varied in their ability to recapitulate the *ex vivo* tumor ultrastructure, with emboli displaying the closest resemblance in both nuclear and LD morphology. We note that these comparisons are simplistic in that they overlook the spatial variance of organelle shapes. To address this, we calculated nuclear aspect ratios and remapped these values onto the original segmentation using an LUT heat map **(Supp Fig S3, bottom)**. This revealed an unexpected gradient: nuclei with high aspect ratios (flatter or oblong; dark red in heat map) were enriched at the suspension periphery, whereas nuclei with low aspect ratios (almost round; dark blue in heat map) were enriched in the center, with measurements similar to tumors. Future experiments will determine if radial gradients in cell surface markers or other functional metrics, if any, accompany these ultrastructural variations. Large-area volume EM imaging combined with automated, high-quality segmentation pipelines can thus uncover unexpected spatial insights in a variety of model systems. When combined with additional data (Balamurugan et al., 2026), our ultrastructural study supports emboli as a compelling i*n vitro* surrogate to study this model of breast cancer.

**Figure 4.**
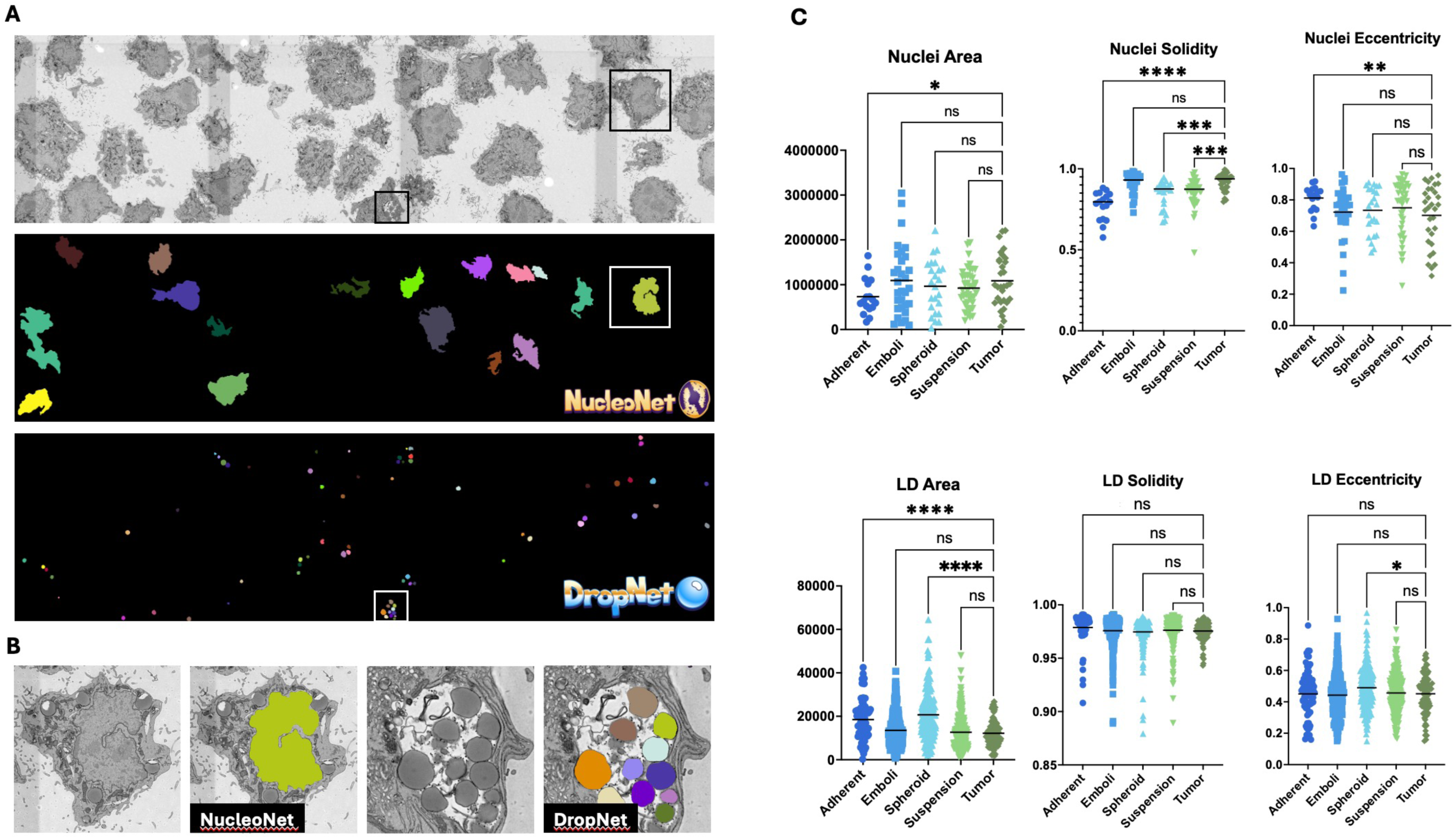
Segmentation and statistically robust shape comparisons enabled by *NucleoNet* and *DropNet*. (A) Adherent SUM149 cells imaged by AT at 5 nm pixel size (top) were segmented by *NucleoNet* and *DropNet* to reveal instances of nuclei (middle) and lipid droplets (bottom) in 2D. Horizontal Field Width of screenshots is 260 μm. (B) Zoomed in paired views of EM data (left) and *NucleoNet* or *DropNet* (right) model predictions of individual cells boxed in A. (C) Comparison of selected 2D shape parameters across 2D (adherent) and 3D (emboli, spheroid, suspension) cell culture models versus tumor tissue. The top row presents nuclear measurements (adherent, n = 18; emboli, n = 32; spheroid, n = 25; suspension, n = 43; tumor, n = 32), and the bottom row shows lipid droplet (LD) measurements (adherent, n = 73; emboli, n = 319; spheroid, n = 187; suspension, n = 258; tumor, n = 91) of area, solidity, and eccentricity. Statistical significance was assessed by parametric unpaired t-tests with Welch’s correction; ns = not significant, * P < 0.05, ** P < 0.01, *** P < 0.001, **** P < 0.0001. Center lines represent median values.

## Discussion

Using a combination of publicly shared volume EM (vEM) data and a crowdsourced annotation pipeline tailored for EM data, we have developed two new deep learning models, *NucleoNet* and *DropNet.* These models generate accurate and precise instance segmentation predictions for nuclei and LDs, respectively. To the best of our knowledge, both are the first freely available models dedicated to these organelles in EM images and easily deployed via a GUI. This work thus addresses a significant gap in existing EM segmentation solutions and expands organelle analysis beyond mitochondria. Both models are available in the latest release of our napari plugin, *empanada v1.2*, allowing point-and-click operations – not just for running inference, but also for fine-tuning, proofreading, and other tasks. Both models show good results with diverse benchmarks, which we also share to facilitate future research and testing. Finally, we show the utility of *NucleoNet* and *DropNet* by using the models to generate large numbers of instance segmentations, allowing quantitative comparisons between *in vitro* tumor models and actual *in vivo* tumors. The following sections describe several pertinent aspects of this work and its relevance to the broader field.

*NucleoNet* accelerates segmentation of nuclei across diverse cell types, and we show good results in our varied benchmarks That said, we acknowledge its limitations. Key among them is the model’s restriction to non-dividing cells, limitation to room temperature vEM, and the inability of the current version to classify sub-regions into heterochromatin, euchromatin and nucleoli. During assembly of the corpus of image patches for model training, we were unable to obtain enough instances of dividing cells. Unlike mitochondria, cells typically have only one nucleus, with some notable exceptions like multinucleated cells arising from cell fusion or neoplasic transformation (Hazra et al., 2023; Lee et al., 2025; Liu et al., 2024; Mirzayans et al., 2018). Further, collated images patches were highly variant in terms of division stage, sample preparation, and imaging conditions. We theorize that this variation, coupled with the hazy staining of DNA and absence of nuclear membranes due to envelope breakdown, contributed to a very weak class which adversely affected model performance. For these reasons we excluded patches from dividing nuclei from training. Mammalian cell nuclei are also relatively large, meaning that training patches of a given size and resolution typically only contained a fraction of a single nucleus. This reduced the number of nuclear instances in any given number of patches, resulting in a modest training dataset compared to *MitoNet*. This paucity was also the reason we chose not to extend training to cryoET images. We also decided not to create a separate class for nuclear sub-structures as they were sparsely represented in the dataset, and student annotators could not reliably annotate nucleoli, euchromatin or heterochromatin; so for practical reasons we focused on nuclei as a whole. Nevertheless, *NucleoNet* can be leveraged to create masks for large numbers of nuclei, facilitating segmentation and/or model training on sub-nuclear features in future studies. Similarly, *DropNet* was not trained on extremely large LDs, which are sometimes observed in pathological cases (Gluchowski et al., 2017; Sugawara et al., 2021). The extracted patches of these droplets often appeared as a uniform gray, with contrast and texture similar to the embedding resin, especially when missing the context of the limiting membrane. Although most LDs are round and have smooth surfaces (i.e. high roundness and circularity), many aspherical and ruffled droplets were observed and included in the training dataset. It is possible that certain intracellular compartments like endosomes that are stained an even gray could be segmented by the model.

Much of the poor performance of DL models in pathological samples, irrespective of organelle segmented, can be ascribed to atypical and highly variable structures that are rarely or never represented in the training corpus (Aswath et al., 2023; Karabağ et al., 2023). This promises to be a fertile area for future research, and we anticipate that the curation of massive training datasets, multimodal data integration, and creation of foundational models and will push the EM field forward.

While these models yield reasonable outputs and provide strong initial segmentations, they are not perfect; we urge users to consider these as “base models” to be fine-tuned to their own datasets for better results (fine-tuning can be performed easily in *empanada*). We encountered some false positive and false negative errors, some split and merge errors, and in the case of *NucleoNet*, some “holey” predictions (high confidence at the nuclear periphery near the nuclear membrane but reduced confidence in the nuclear centers). A larger and more diverse pre-training and training dataset, especially one including convoluted nuclei and crowded LDs, would likely improve performance. For “holey” nuclear predictions, the limitations stem from both the inherent weakness of the PDL architecture and small patch size in training, which reduce predictive confidence in patches lacking membranes or distinct edges; conversely confidence is higher when a nuclear membrane is present. A suggested solution is to downsample high resolution images to capture some nuclear membrane within the larger resulting patches. We consequently observe significantly better results, but we caution that overaggressive downsampling can obscure fine features. Our models perform optimally in the 15 - 40 nm pixel range in line with the majority of training data resolutions, as described previously (Conrad and Narayan, 2023). Larger and more sophisticated models, which can be tested against our nuclear EM benchmarks, will alleviate this issue in the future.

Looking ahead, newer approaches such as vision transformers (Dosovitskiy et al., 2020; Hatamizadeh et al., 2022; Hörst et al., 2024) and diffusion models (Aswath et al., 2023; Lu et al., 2024) show great promise for segmenting large and 3D image data. Likewise, developments in multi-class segmentation, where all cellular organelles are segmented simultaneously with a single model, have already spawned a challenge in the field, and we ourselves leveraged this resource to test an nnU-Net against *NucleoNet* (https://cellmapchallenge.janelia.org/). The rapid generation of large numbers of instance segmentation data with minimal manual interventions will enable morphological measurements with large sample sizes, enabling robust statistical analyses of EM data and supporting truly quantitative ultrastructural studies in cell biology. We anticipate that *NucleoNet* and *DropNet* will accelerate biological discovery by enabling robust, automated and high-throughput visualization and quantitation of nuclear and LD morphology directly from EM data; future models for other organelles will also help. Another intriguing use-case is for high-throughput or large -volume correlative microscopy, where fluorescence images of stained nuclei can be registered to automatically segmented nuclei using point-cloud based approaches, similar to recently published work(Krentzel et al., 2025). We are realistic that future solutions from AI breakthroughs will supersede our current results. Yet we expect our models, especially when paired with the intuitive GUI of our napari plugin *empanada*, will find immediate utility within the bioimaging community.

## Materials and Methods

### EM Datasets

Training datasets included three room temperature EM modalities (no cryoEM/ET data was included): Focused Ion Beam Scanning Electron Microscopy (FIB-SEM), Array Tomography (AT) and Transmission Electron Microscopy (TEM). Images were sourced from a wide range of eukaryotic biological samples, including mammalian and non-mammalian cells and model organisms; tissues, isolated cells, cell lines; prepared by both conventional fixation and high-pressure freezing. Pixel sampling ranged from 10 nm to 45 nm/pixel. The final training dataset for the nucleus model consisted of [1024 x 1024 x 1] sized patches from 31,082 images, comprising 16,072 FIB-SEM images, 2,130 TEM images, 10,380 AT images and 2,500 serial block-face SEM (SBF-SEM) modalities. Of these, 17,746 images were sourced internally from FNL/NCI archives and 13,336 were sourced externally from publicly available repositories, including https://openorganelle.janelia.org/, http://www.nanotomy.org/, https://www.ebi.ac.uk/empiar/, https://www.microns-explorer.org/ and https://www.ebi.ac.uk/empiar/EMPIAR-10365/. This dataset contained 25,573 images from tissue samples and 5509 images from cell cultures. For the lipid droplet (LD) model, the training set comprised 2,042 images which included 1,792 images with positive instances of LDs and 250 images with no LDs. During collection, all datasets were standardized to 8-bit unsigned volumes/images, binned to normalize resolution, transformed to isotropic volumes if required, and finally stored as ZARR file volumes, from which patches were extracted, as below.

### Image Preprocessing

Patchification: 3D volumes were sliced along orthogonal views, and a sliding window was used on 2D images to generate non-overlapping 1024 x 1024 2D images, resulting in ∼2.5 million patches from internal datasets and ∼4.8 million patches from external datasets.

Deduplication: To remove redundant images, difference hashing with a hash size of 8 was used. Duplicates were identified by calculating the Hamming distance between hashes. Patches with a distance below a threshold of 12, a value determined through visual inspection, were discarded. This reduced the patch count to approximately 242,000 for internal and 471,000 for external sources.

Filtering: Non-informative patches, such as those containing resin artifacts or excessive noise, were excluded using a classifier adapted from the CEM500K dataset Specifically, an ImageNet-pretrained ResNet34 classification model was trained on 12,000 images manually classified as informative and non-informative. After filtration, the final dataset consisted of approximately 148,000 internal and 297,000 external patches.

Classification: Given the enormous number of patches to be annotated and limited annotators, a YOLO-based strategy was implemented to efficiently select heterogenous and “difficult” patches for downstream crowdsourced annotation. First, a simple Jupyter notebook was used to manually classify organelle-rich images, creating seed batches for annotation. Images presented to the human classifier were chosen by metadata bucketing with these categories: image modality, pixel spacing, biological context and cell/tissue. Images were randomly selected from each bucket and presented to a human annotator to create batches of roughly 500 images, which were uploaded to Zooniverse for annotation. A YOLO11n-seg model (https://docs.ultralytics.com/tasks/segment/#models) network was trained on segmentations from these initial batches and then used for prediction across the final filtered dataset pool. Patches where the detection confidence threshold was low were chosen to be shown to a human annotator via an interactive Jupyter notebook to classify an image as worthy of annotation. This iterative selection enriched the image pool with challenging patches likely to provide valuable information for model training, while avoiding patches too difficult that annotators would struggle to create reliable ground truth (GT) labels. The YOLO network identified 42,467 internal and 28,165 external images as “difficult”. From these, a random subset of ∼25,000 images, uniformly sampled across different metadata categories, was manually classified as to-be (or not-to-be) annotated, and approximately 500 images at a time were released for Zooniverse annotation.

### Nucleus Image Annotation via Zooniverse

Data annotation patches were hosted on the web-based citizen science platform Zooniverse (www.zooniverse.org) for annotation, and a cohort of high-school students added as annotators to the project. The project was configured with a retirement limit of five annotations per image, and roughly 500 images per batch were uploaded weekly. Preprocessed EM images were uploaded to the Zooniverse platform using an uploader module which utilizes the Panoptes Client API. The annotation interface for this project was designed with specific instructions and tools: a polygon drawing tool with adjustable vertex placement, reference images and a field guide showing example nuclei annotations, and a tutorial and help button for additional guidance and FAQs (Spiers et al., 2021b). Student annotators were tasked with drawing polygons around nuclei in the images displayed outlining the nuclear membranes as accurately as possible while disregarding all other organelles. Images containing multiple nuclei were included, with the expectation that each nucleus would be annotated. On rare occasions when nuclear “donuts” were present, students were instructed to segment it as a “keyhole” due to limitations with the drawing tool, and we relied on the downstream consensus algorithm to reveal the donut hole. Each image was randomly presented to five students, and an expert annotator also contributed by annotating approximately fifty images. At the end of every batch (typically a week), the Zooniverse platform reported the number of images annotated by each participant. Annotation accuracy was assessed by comparing each student’s output to the expert’s annotations, which served as the GT for scoring. Student annotation output and accuracy were combined to yield a score, and a leaderboard was generated. We found that this incentivized the students to perform accurate segmentations at a reasonable rate, and we were able to identify poor performers for individualized help.

### Lipid Droplet Image Annotation

DropNet was trained using a diverse collection of TEM, SEM, and FIB-SEM images, totaling 25 datasets that included breast cancer cells (SUM149), mouse biopsies (kidney, liver, and pancreas), and *macrostomum Lignano*. These datasets varied in resolution from 5 nm – 25 nm in the xy plane. The final training dataset consisted of 2,042 images (size: 512 x 512 x 1 pixels), with 1,792 containing LDs and 250 without. For annotation, four true positive classes were defined via manual segmentation in *empanada-napari*: LDs with a dark membrane, LD without a defined membrane, LD with clear holes stemming from preparation limitations, and LD with irregular staining and shapes. The true negative class included: alpha, beta, gamma pancreatic islet cell images, and images without LDs. Training images with LDs contained between one and 50 instances per image.

### Model Training with empanada

The *empanada* library, which powers the *empanada* napari plugin, was used to train nuclei (*NucleoNet*) and lipid droplet (*DropNet*) segmentation models, ensuring compatibility with the napari plugin and enabling the use of CEM1.5M pretrained weights (Conrad and Narayan, 2023) for improved accuracy and domain generalization. The *empanada* library config file-based setup made hyperparameter testing efficient and streamlined. Training employed the *empanada* framework with an extensive pipeline using the Albumentations library (Buslaev et al., 2020), including random 90-degree rotations, random flips, random elastic transform deformations (a = 120, s = 120 x 0.05, a_affine_ = 120 x 0.03), brightness and contrast adjustments (brightness limit = 0.2, contrast limit = 0.2), Gaussian noise (variation limit = (0, 25.0)), and Gaussian blur (blur limit = (1, 3)), all to boost generalization. A composite loss function was used: cross-entropy for sematic segmentation, focal loss for center prediction (a = 0.25, g = 2.0), and L1 loss for offset prediction. The total loss was computed as 1.0 x semantic_loss + 200.0 x center_loss +0.01 x offset_loss. Optimization was conducted with the Adam optimizer (learning rate 1 x 10^-4^, weight decay 1 x 10^-4^), applying weight decay only to weights (not biases), and excluding biases and Batch Norm layers. A polynomial learning rate decay (power = 0.9), batch size of 8 per GPU, and gradient accumulation for an effective batch size of 16 were used. The training strategy involved transfer learning, initializing encoders with CEM1.5M weights and progressively unfreezing encoder layers: decoder-only for 10 epochs, then unfreezing encoder stages (stage4 and stage3) over the next 20 epochs, followed by all layers for the final 70 epochs. Training ran on 4 NVIDIA A100 GPUs with distributed parallelism, taking 24 hours for *NucleoNet* and 4 hours for *DropNet*. Model validation included monitoring mean IoU and average precision (AP) at IoU thresholds of 0.5, 0.75, and 0.9 every 5 epochs, with early stopping with patience after 15 epochs based on validation of a 0.75 AP. empanada-napari download and documentation, including tutorials, is available at https://empanada.readthedocs.io/en/latest/index.html.

### Model comparison with CellSAM and nn-Unet

To assess zero-shot generalization performance, we directly applied CellSAM (Marks et al., 2025) to our benchmarks using the publicly available cellsam_general (v1.2) model without any fine-tuning or parameter adaptation. Segmentation was performed in 2D slice-wise mode using the WSI inference strategy implemented in CellSAM, with a block size of 1024 pixels, 512-pixel overlap, IoU depth of 256, IoU threshold of 0.55, and bounding-box threshold of 0.5. Predictions were generated independently for each slice of the volume. Small instances with an area smaller than 1,200 pixels were removed during post-processing. In the case of nnU-net (Isensee et al., 2020a), we specifically trained the model on 21 CellMap electron microscopy volumes (https://cellmapchallenge.janelia.org/) spanning 16 nm, 32 nm, and 64 nm resolutions, covering multiple tissue types including kidney, liver, cardiac tissue, skeletal muscle, and glomerulus. All nuclear annotations were converted into a binary format, and the final model was trained on 2D slices at 32 nm resolution using the standard nnU-Net 2D U-Net configuration. Model training required approximately 8 hours on a single NVIDIA A100 GPU. In both cases, during inference, the modes were applied slice-wise to generate nuclear probability maps, followed by watershed-based post-processing to obtain instance segmentations. A minimum distance of 150 pixels between local maxima was used, and objects with an area smaller than 1,000 pixels were removed. In both cases, inference was conducted on the same Linux-based HPC node equipped with dual Intel(R) Xeon(R) Platinum 8362 CPUs (64 cores total), 256 GB system memory, and NVIDIA A100 GPUs. Processing three benchmark volumes required approximately 50 minutes total wall-clock time for CellSAM, and roughly 15 minutes for nn-Unet including watershed.

### Cell Culture

SUM149 breast cancer cells (Asterand Bioscience) were cultured in Ham’s F-12 media (GIBCO #31765092) supplemented with 10% FBS, 5 ug/ml hydrocortisone (Sigma #H-0135), 1 ug/ml insulin (Sigma #I-0516), 100 units/ml penicillin, 100 ug/ml streptomycin, and 2.25% PEG8000 (Sigma #202452) at 37°C with 5% CO_2_ for adherent, emboli, and suspension growth types. For spheroid growth, SUM149 cells were cultured in Mammocult media (StemCell Technologies #05620) supplemented with 2 ug/ml heparin (StemCell Technologies #07980) and 0.44 ug/ml hydrocortisone. For emboli culture, ultra-low attachment (ULA) multi-well plates (Corning 6-well; #3471 or 24-well; #3473) were used and performed as previously described (Balamurugan et al., 2025; Lehman et al., 2013). Three-dimensional cell cultures (emboli, suspension, and spheroid) were harvested by centrifugation at 500 rpm (30 s), washed with PBS, treated with TrpLExpress (GIBCO #12604-013) for 10 minutes with intermittent agitation, and neutralized with cell culture media. Adherent SUM149 cells were isolated by trypsinization. Tumors were isolated and fixed as described in (Balamurugan et al., 2025).

### Sample Preparation and Imaging

After isolation, all cell growth types were washed with PBS and fixed with 4% formaldehyde/2% glutaraldehyde in 0.1 M sodium cacodylate buffer for 5 hours at room temperature (RT) with occasional rocking to prevent clumping. Samples were centrifuged (500 rpm; 30 s) and the supernatant decanted to remove fixative. Samples were resuspended in 0.1 M sodium cacodylate buffer and centrifuged again. Cells were suspended in the same buffer again and stored at 4°C. Fixed adherent cells were pelleted and embedded in 1% low-melt agarose and fixed 3D cells were left free-floating. Subsequent EM sample preparation followed the same protocol presented in Balamurugan et al., 2025. Polymerized resin blocks were trimmed, and 100 nm sections were collected on ITO coverslips utilizing a Leica ARTOS ultramicrotome. Large-areas of resin sections on ITO coverslips were imaged at 5 nm pixels in 8k x 8k sized tiles in a GeminiSEM 450 with a four-quadrant backscatter detector (aBSD, Zeiss) with a 3.5 kV beam current, 2 kV beam deceleration, and 800 pA probe current using ATLAS 5 Array Tomography software (Fibics). Final image tiles were stitched through the ATLAS 5 software.

### Model Segmentation and Image Analysis

Images were cropped to yield comparable cell numbers across the samples (Adherent, Emboli, Spheroid, Suspension, and Tumor). These images were imported into napari (Chiu et al., 2022) where segmentation was performed using the 2D inference module in the *empanada* plugin with appropriate DL models: nuclei (NucleoNet_base_v1) and LDs (DropNet_base_v1). Errors in model predictions were corrected in *empanada* utilizing the ‘merge labels’, ‘split labels’, and ‘morph labels’ modules. To accelerate these corrections, the ‘create tiles’ module was used to generate smaller image tiles. Following correction, images and their corresponding segmentations were reassembled using the ‘merge tiles’ module. Segmentations that contacted the image border were removed using the ‘filter labels’ module prior to any measurements. Morphometric quantitation of the cell nuclei and lipid droplet segmentations was performed using the clusters-plotter plugin, which applies region props from scikit-image (Van Der Walt et al., 2014; Zigutyte et al., 2025). Graphs were generated and statistical analyses performed using Prism (Ver. 10.5.0 for Mac). Differences between growth conditions were analyzed using a nonparametric unpaired one-way ANOVA Kruskal-Wallis test with a p-value of 0.05 being significant.

## Acknowledgments

The crowdsourced annotated portion of the training dataset used for NucleoNet was generated by a cohort of high school students from the Frederick County Public School system (FCPS), MD, USA. We thank the student participants and teacher advisors. We thank Mariam A Demir from Science and Technology Facilities Council (STFC), UKRI, for empanada support. This project has been funded in whole or in part with Intramural funding or Federal funds from the National Cancer Institute, National Institutes of Health, under Contract No. 75N91019D00024. The content of this publication does not necessarily reflect the views or policies of the Department of Health and Human Services, nor does mention of trade names, commercial products, or organizations imply endorsement by the U.S. Government.

## Figure Legends

**Figure S1. Nuclei examples used in *NucleoNet* model training.**

Randomly chosen patches from datasets used for crowdsourced annotations. Nuclei included were from various sources like nuclei from adherent or nonadherent cell from *in vitro* experiments, as well as simple and complex nuclei from *in vivo* tissue samples.

**Figure S2: Lipid droplet classes examples used in *DropNet* model training.**

Three examples each of (A) lipid droplets (LD) with a slightly variable staining and an obvious dark outline, (B) evenly stained LDs without a defined border, (C) LDs with “holes” resulting from sample preparation limitations, and (D) LDs with irregular shapes and other irregularities like pronounced variable staining.

**Figure S3. Growth of cell culture models and spatial variation in nuclear morphology.**

A. Brightfield images of 2D (adherent, top left) and 3D cell culture models (suspension, top right; emboli, bottom left; spheroid, bottom right). Scale bars are 400 μm.
B. Cross section of a 3D spheroid imaged by AT with nuclei segmented by *NucleoNet* (left), enabling extraction of shape metrics such as aspect ratio. Aspect ratio values for individual nuclei are projected back onto the segmentation and visualized as a LUT heat map (right), with dark red indicating nuclei with high aspect ratios (closer to 1; flatter or more elongated shape) and dark blue indicating nuclei with low aspect ratios (closer to 0; more circular in shape). Horizonal Field Widths for snapshots are 400 μm.

## Notes

https://empanada.readthedocs.io/en/latest/index.html

### Competing Interest Statement

The authors have declared no competing interest.

