## Supplementary material for "NucleoNet and DropNet: Generalist deep learning models for instance segmentation of nuclei and lipid droplets from electron microscopy images": Table 1

| Image | F-1@IoU0.5 | F-1@IoU0.75 | TP_fraction | FP_fraction | FN_fraction |
|---|---|---|---|---|---|
| Zebrafish | **0.78571** | 0.61905 | 0.64706 | 0.15686 | 0.19608 |
| M.lignano | **0.81818** | 0.66667 | 0.69231 | 0.23077 | 0.07692 |
| Lymphoid Organ | **0.63333** | 0.53333 | 0.46341 | 0.31707 | 0.21951 |
| Spleen | **0.7** | 0.6 | 0.53846 | 0.23077 | 0.23077 |
| Liver Cancer | **0.7451** | 0.70588 | 0.59375 | 0.21875 | 0.1875 |
