## Supplementary material for "NucleoNet and DropNet: Generalist deep learning models for instance segmentation of nuclei and lipid droplets from electron microscopy images": Table 2

| **Pixel-Level Metrics (NucleoNet PRED)** | **Pixel-Level Metrics (CORRECTED)** | **Instance-Level Metrics (NucleoNet PRED)** | **Instance-Level Metrics (CORRECTED)** |
|---|---|---|---|
| Overall Accuracy: 0.9956 | Overall Accuracy: 0.9935 | True Positives (TP): 296 | True Positives (TP): 296 |
| Label 0 (background): 0.9976 | Label 0 (background): 0.9983 | False Positives (FP): 218 | **False Positives (FP): 20** |
| Label 1 (nuclei): 0.9637 | Label 1 (nuclei): 0.9151 | False Negatives (FN): 51 | False Negatives (FN): 51 |
| | | Precision: 0.5759 | **Precision: 0.9367** |
| | | Recall: 0.8530 | **Recall: 0.8530** |
| | | F1 Score: 0.6876 | F1 Score: **0.8929** |
| Mean IoU (Pixel-Level): 0.9610 | Mean IoU (Pixel-Level): 0.9421 | Mean Instance IoU: **0.8732** | Mean Instance IoU: 0.8679 |
| Mean Dice (Pixel-Level): 0.9798 | Mean Dice (Pixel-Level): 0.9695 | Mean Instance Dice: 0.9252 | Mean Instance Dice: 0.9222 |
