## Supplementary material for "NucleoNet and DropNet: Generalist deep learning models for instance segmentation of nuclei and lipid droplets from electron microscopy images": Table 3

| image | F-1@0.5 | F-1@0.75 | TP_fraction | FP_fraction | FN_fraction |
|---|---|---|---|---|---|
| Fatty Liver | 0.77573 | 0.77045 | 0.63362 | 0.30172 | 0.06466 |
| Pancreas | 0.93333 | 0.91111 | 0.875 | 0.0625 | 0.0625 |
| Breast Cancer | 0.89 | 0.84 | 0.8018 | 0.13514 | 0.06306 |
