## Supplementary figures and images for "NucleoNet and DropNet: Generalist deep learning models for instance segmentation of nuclei and lipid droplets from electron microscopy images"

### Supplemental Figure 1

#### Fig S1

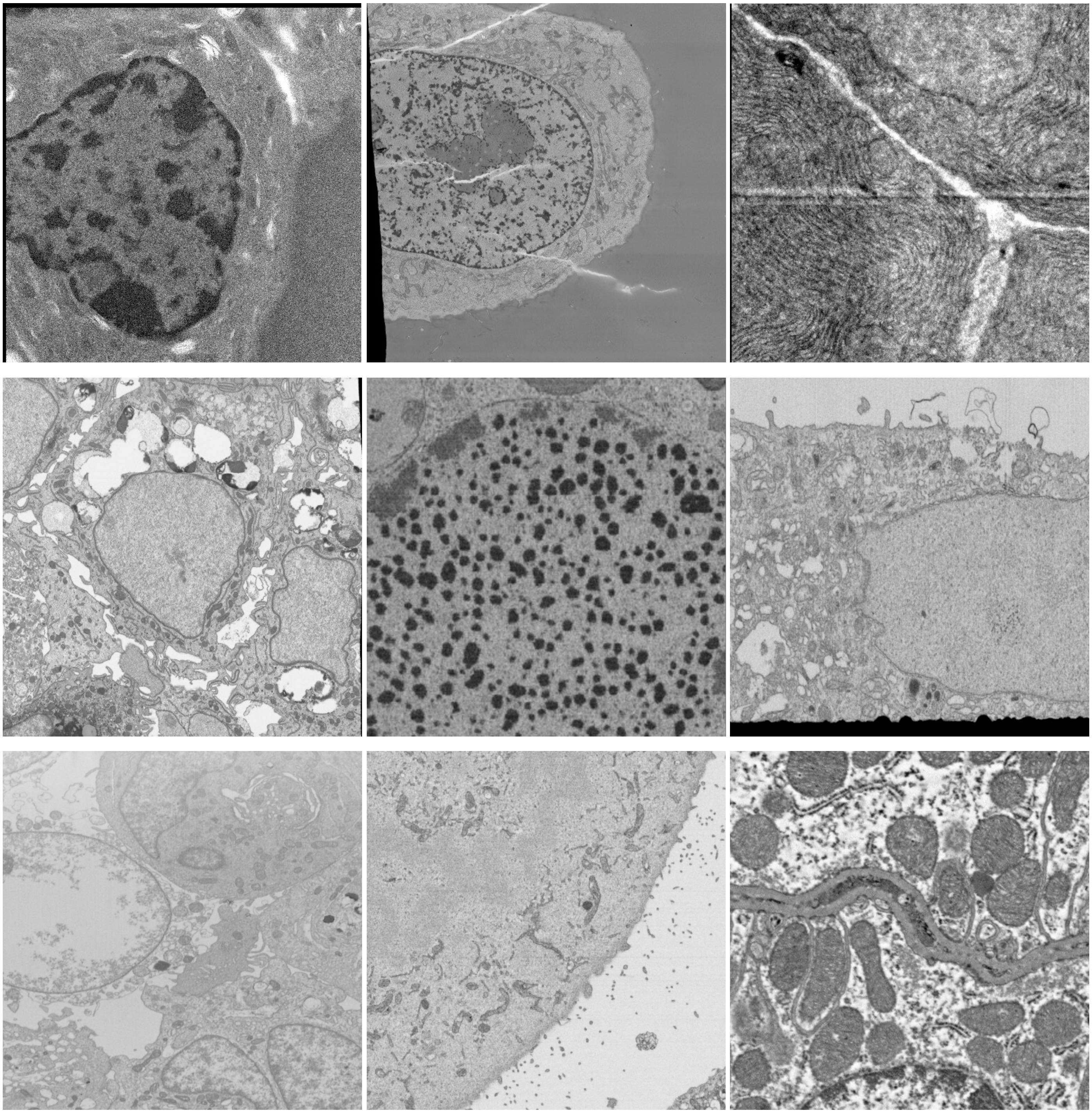

### Supplemental Figure 2

**Fig S2**

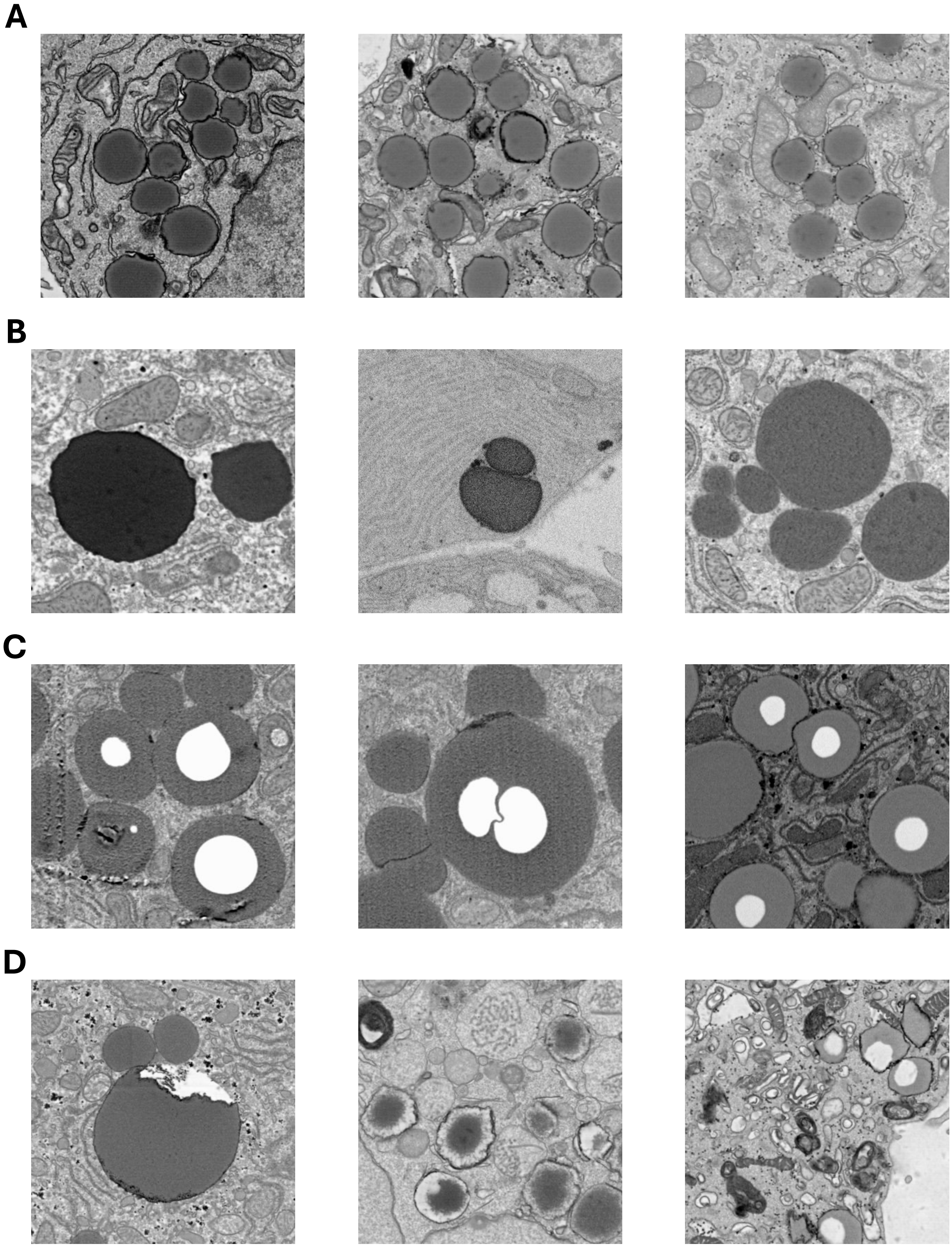

### Supplemental Figure 3

#### Fig S3

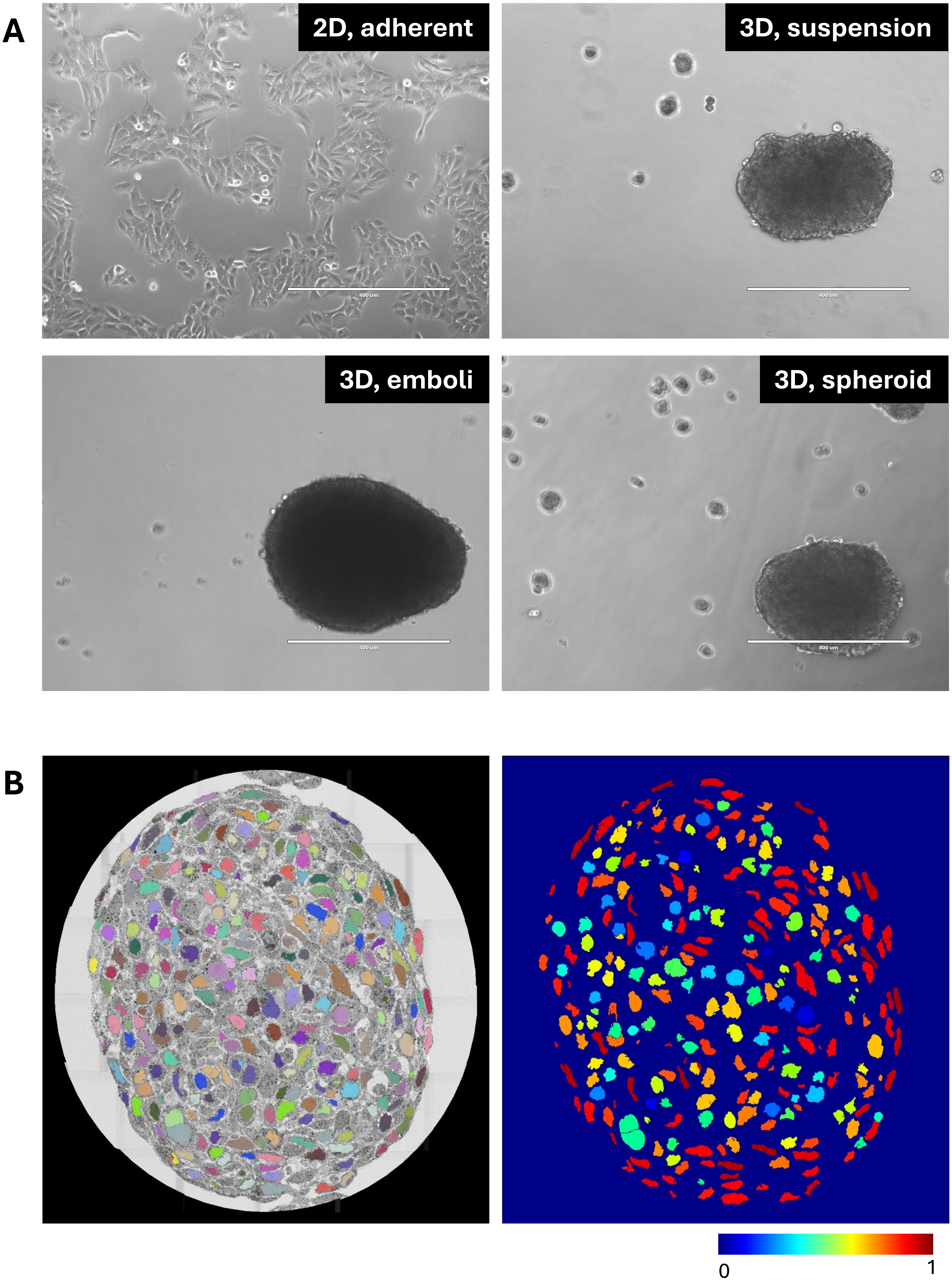
